# Oncogenes and tumor suppressor genes are enriched in stop-loss mutations generating protein extensions

**DOI:** 10.64898/2026.03.12.711331

**Authors:** Lilian M. Boll, Joan Antoni Martorell, Nafiseh Khelghati, Marta E. Camarena, Clara Vianello, Juan Carlos García-Soriano, Eva Santamaría, Idoia Artoleta, Silvia Sáez-Valle, Júlia Perera-Bel, Puri Fortes, M. Mar Albà

## Abstract

Cancer genomes tend to accumulate a large number of mutations, and even rare mutations such as those causing the loss of a stop codon can be observed in a significant fraction of the tumors. Stop-loss mutations extend protein translation into the 3’ untranslated region (3’ UTR), generating altered proteins carrying extra amino acid sequences. These C-terminal extensions can potentially have consequences for tumorigenesis and immune recognition. To investigate the prevalence of stop-loss mutations in cancer, and to identify recurrent mutations with a possible tumor-promoting effect, we have interrogated mutation data from the tumor samples of 20,801 patients. This search has resulted in the annotation of 3,757 stop-loss mutations in 3,249 different protein-coding genes. Around 11% of the mutated genes contain recurrent stop-loss mutations, occurring in more than one patient. The protein extensions created by the mutations tend to be hydrophobic and/or positively charged, and these features are associated with an increased propensity to generate MHC I-bound peptides. We have also found that cancer-related genes contain 37% more stop-loss mutations than non-cancer-related genes, with both oncogenes and tumor suppressor genes showing similar enrichments. Furthermore, three out of the four genes with the highest number of stop-loss recurrences, *PTMA*, *PCDH9 and SOX9*, are cancer-related. In *PTMA*, the gene with the largest number of stop-loss mutations (14 patients), the mutation results in an extension of 9 amino acids. We provide experimental evidence that the mutation is associated with impaired cleavage of thymosin alpha 1, a peptide with immunostimulatory functions that is generated from the N-terminal part of the PTMA protein. The study provides evidence that stop-loss mutations are enriched in cancer-associated genes and constitutes a valuable resource for further studies on the effects of stop-loss mutations in cancer.

## INTRODUCTION

Stop codons (TAA, TAG, and TGA) terminate the ribosomal translation of a coding sequence. A mutation converting a stop codon to a sense codon, referred to as a stop-loss, stop-lost, or non-stop mutation, causes the continuation of translation elongation into the 3’ untranslated region (UTR). The elongation continues up to the next in-frame stop codon or the polyA tail of the mRNA molecule. The latter cases are rare, and their translation is expected to be limited by destabilizing and accelerated degradation of the mRNA molecule and the nascent protein (Klauer and van Hoof 2012; Shibata et al. 2015). Some stop-loss mutations cause heritable diseases; for example, in *CLDN11*, encoding claudin-11, stop-loss variants have been associated with neuronal abnormalities (Riedhammer et al. 2021). In general, C-terminal extension length, hydrophobicity and protein aggregation propensity have been positively linked to increased pathogenicity (Takata et al. 2021; Yoon et al. 2026).

A previous survey of mutations centered in the COSMIC database of cancer genes identified stop-loss mutations in *SMAD4* in several cancer patients (Dhamija et al. 2020). The 40 amino acid protein extension was associated with increased degradation of the protein via a newly created hydrophobic degron sequence. Other identified examples of protein inactivation due to stop-loss mutations were the renal tumor suppressor genes *BAP1*, *PTEN*, and *VHL* (Pal et al. 2025). While the short stop-loss extensions of *PTEN* and *VHL* are thought to affect the proteins’ stability, leading to proteasomal degradation, stop-loss mutations in *BAP1* result in a very long extension that leads to translation inhibition of the mRNA. It was also shown that, in general, protein degradation-promoting extensions tended to have higher hydrophobicity than those that did not have such an effect (Ghosh et al. 2024).

In a previous study, we reported that a high stop-loss mutational burden is significantly associated with a positive response to immunotherapy in bladder cancer (BLCA) patients (Boll et al. 2025). This could imply that some of the extensions could generate peptides that bind to HLA I and are recognized as “foreigner” by the immune system. The potential immunogenic effect of peptides originating from the readthrough of a stop codon is supported by the work of Goodenough and colleagues where aminoglycosides, a drug against premature stop codon mutations, was found to enhance stop codon readthrough, leading to an increased CD8+ T cell response due to MHC I-presented epitopes (Goodenough et al. 2014).

To better understand the impact of stop-loss mutations in cancer, we have gathered mutation data from 20 different cancer patient cohorts (20,801 patients) and analyzed their occurrence in the complete set of 23,231 protein-coding genes. This has increased our initial set of 85 stop-loss mutations in BLCA (Boll et al., 2025) to 3,757 stop-loss mutations in multiple cancer types. Using this data, we have observed a significant enrichment in stop-loss mutations in both tumor suppressor genes and oncogenes. We show that these alterations can have an effect on the properties of the PTMA protein, in which recurrent stop-loss cases are observed. The compendium of stop-loss mutations, and the resulting C-terminal extensions, will contribute to a better understanding of this poorly studied mutation type.

## RESULTS

### A compilation of stop-loss mutations in cancer samples

We compiled 3,757 stop-loss mutations from 20,801 cancer patients from 20 different cohorts (**Figure 1A**)(**Table S1**). The mutation data was obtained from several large pan-cancer projects - The Cancer Genome Atlas (TCGA)(Weinstein et al. 2013), the International Cancer Genome Consortium (ICGC)(International Cancer Genome Consortium 2010), the Hartwig Medical Foundation (HMF, https://www.hartwigmedicalfoundation.nl), the Cancer Genome Characterization Initiative (CGCI)(https://ega-archive.org/studies/phs000235) and the Clinical Proteomic Tumor Analysis Consortium (CPTAC)(Edwards et al. 2015). Additionally, we obtained data from smaller size studies dedicated to particular cancer types, including bladder cancer (Snyder et al. 2017; Mariathasan et al. 2018; Miao et al. 2018; Boll et al. 2023), lung cancer (Roper et al. 2023), multiple myeloma (Skerget et al. 2024), hepatocellular carcinoma (Löffler et al. 2019), melanoma (Bassani-Sternberg et al. 2016; Chong et al. 2020) and prostate cancer (Quigley et al. 2018)(**Table S2**).

**Figure 1.**
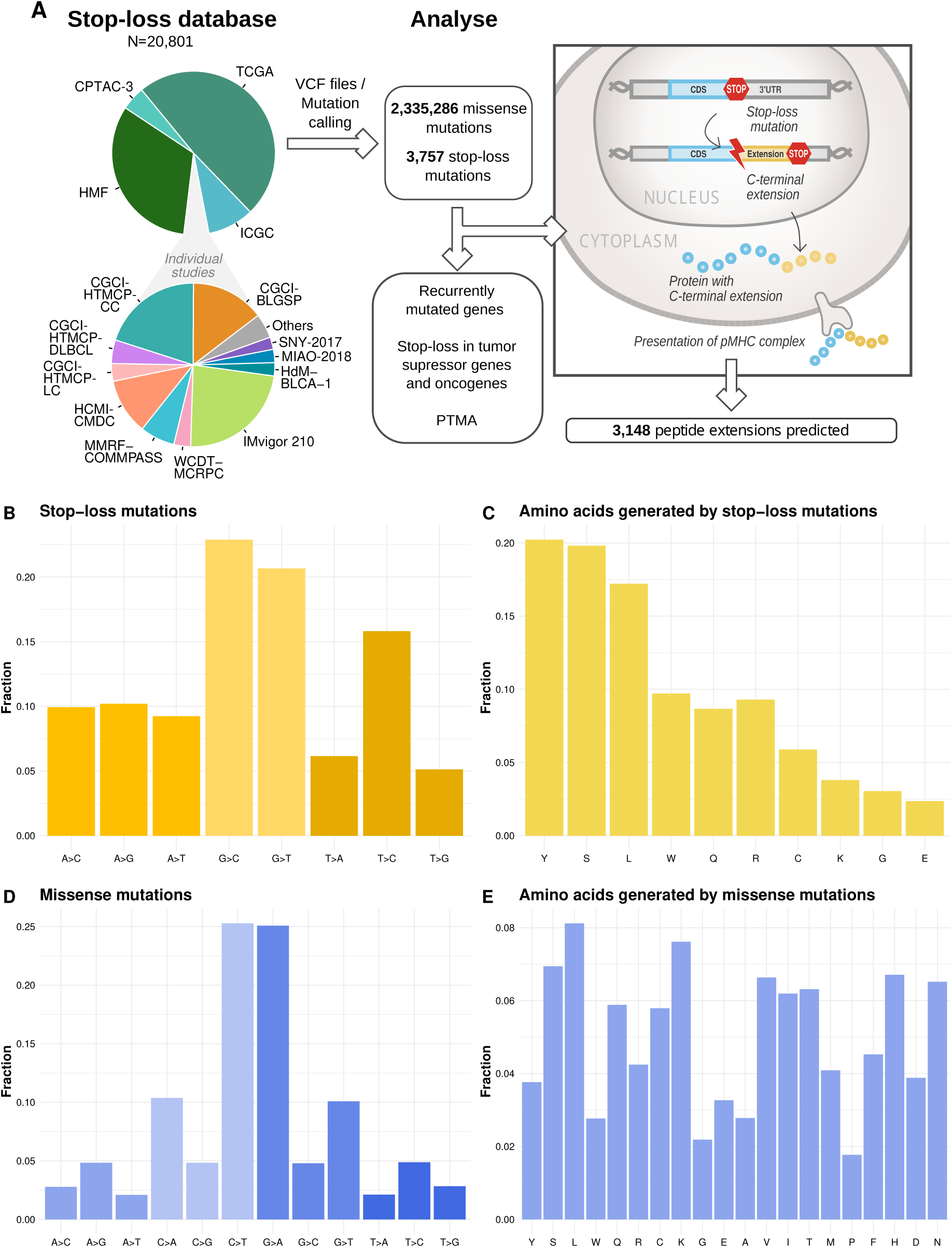
Compilation of stop-loss mutations and missense mutations. **A.** Source of the data and summary of the bioinformatics analysis pipeline. Mutation data from 20,801 patients was obtained from different cancer cohorts. TCGA: Cancer Genome Atlas; HMF: Hartwig Medical Foundation; ICGC:

The vast majority of the stop-loss mutations were single nucleotide variants (3,726 stop-loss mutations, 99%). The mutations affected 3,249 different protein-coding genes out of the 23,231 analyzed (14%). Overall, we identified stop-loss mutations in the tumors from 2,533 patients out of the 20,801 analyzed (12.18%).

The most common mutations in stop codons leading to a sense codon were G>C, G>T and T>C (**Figure 1B**)(**Table S3**). Only 10 out of 20 amino acids can be generated by mutations in the stop codons (**Figure S1**). Of them, the most abundant ones were tyrosine (Y) and serine (S), followed by leucine (L)(**Figure 1C**). For comparison, the number of missense mutations in the same set of samples was 2,335,286. The most common missense mutations were C>T and G>A (**Figure 1D**), and the most abundant amino acids generated by missense mutations were leucine (L) and arginine (R)(**Figure 1E**).

International Cancer Genome Consortium; CPTAC-3: Clinical Proteomic Tumor Analysis Consortium phase 3. **B.** Mutation frequencies for single nucleotide variants corresponding to stop-loss mutations in the complete dataset. **C.** Frequency of the different amino acids generated by stop-loss mutations. **D.** Mutation frequencies for single nucleotide variants corresponding to missense mutations in the complete dataset. **E.** Frequency of the different amino acids generated by missense mutations.

### Stop-loss mutations tend to generate hydrophobic and/or positively charged extensions

We computed a total of 3,148 extensions, of which 2,808 were unique, resulting from the stop-loss mutations (**Table S1**). Cases with no downstream stop codon found in the transcript were discarded (219 transcripts). The median length of the extensions was 19 amino acids, with 95% being shorter than 82 amino acids (**Figure 2A)(Table S4)**.

**Figure 2.**
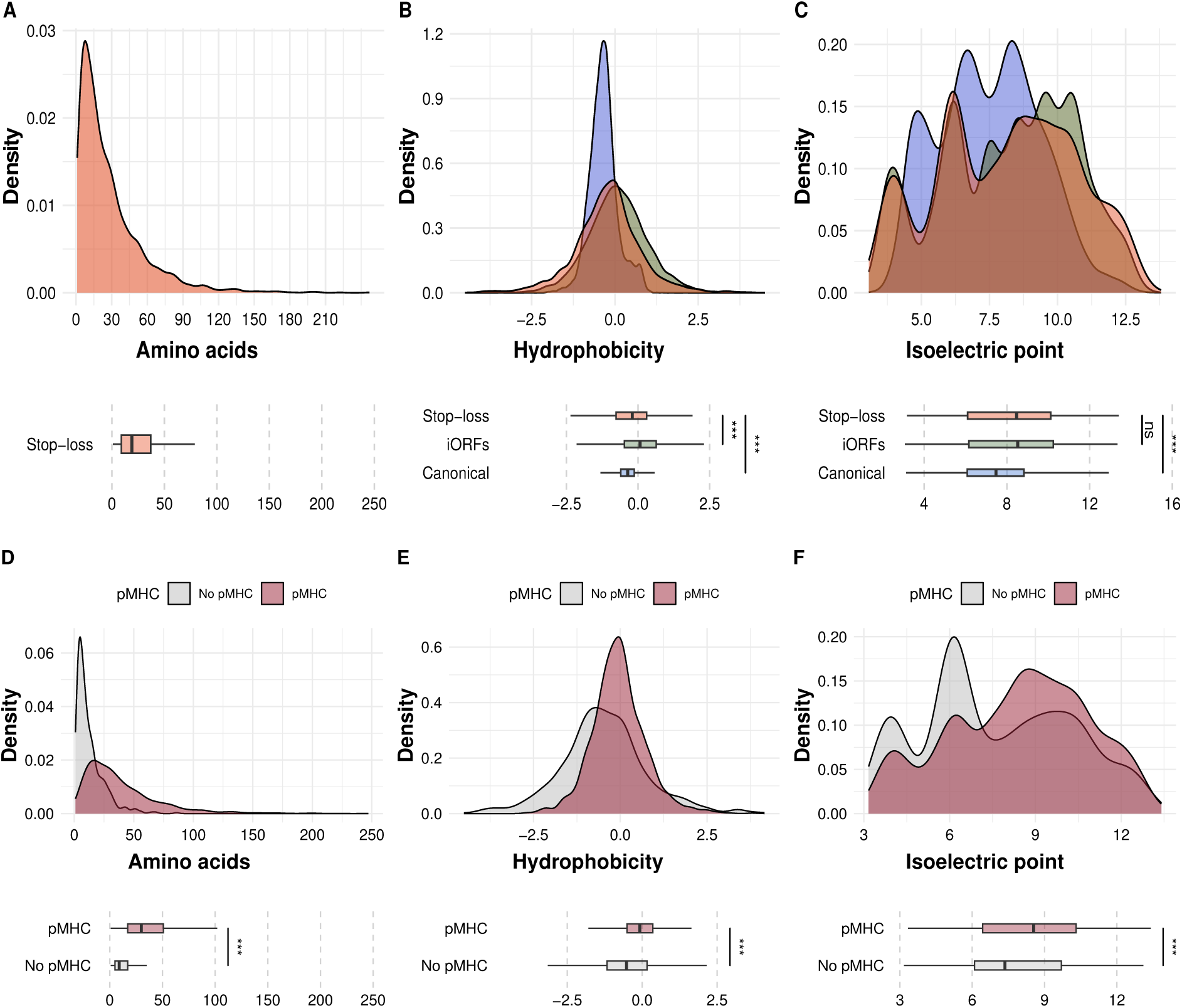
Properties of protein extensions from stop-loss mutations in cancer. **A.** Amino acid length of predicted unique peptide extension resulting from stop-loss mutations (N=2,808). **B.** Hydrophobicity in stop-loss extensions, canonical peptides, and intronic open reading frames (iORFs). Hydrophobicity was computed using the KyteDooLittle scale. **C.** Isoelectric point in the stop-loss extensions, canonical peptides, and iORFs. The isoelectric point was calculated using the EMBOSS pK scale. **D**. Extensions containing at least one predicted strong MHC I binder (pMHC) tend to be longer than the rest. **E**. pMHC extensions tend to be more hydrophobic than the rest (p-value=1.06e-32). **F.** pMHC extensions tend to have a higher isoelectric point (iP) than the rest (p-value = 8.28e-15). All p-values obtained from Wilcoxon-Mann-Whitney test with Benjamini-Hochberg multiple testing correction. Significance levels indicated in the figure are for comparisons of stop-loss extensions versus canonical and stop-loss extensions versus iORFs. Statistical significance is indicated as follows: *** p-value < 10^−14^.

We next compared the hydrophobicity of the protein extensions to those of canonical protein sequences and *in silico* translated intronic regions (iORFs). The latter sequences were taken as a proxy for non-coding sequences. The stop-loss extensions were significantly more hydrophobic than canonical protein sequences, although less than iORFs (Wilcoxon-Mann-Whitney test p-value < 2.22e-16 in both comparisons)(**Figure 2B**). We also found that they tended to have a significantly higher isoelectric point (IP), indicating a more positive charge, than canonical proteins (Wilcoxon-Mann-Whitney test p-value < 2.22e-16) (**Figure 2C**). In this case, no differences with iORFs were observed. One extreme case of a positively charged extension due to a stop-loss mutation was detected in the cyclin encoding mRNA *CCNT1*; the extension contained 8 lysines and 2 arginines out of a total of 12 amino acids. In summary, the extensions have clear compositional biases, which can be essentially explained by the non-coding nature of the sequences encoding them.

### Hydrophobicity and positive charge favor peptide presentation by MHC complexes

The presentation of peptides derived from cancer-specific protein extensions by MHC I and II molecules might enhance the recognition of the tumor by the immune system. This would be consistent with the observation that a high stop-loss mutational burden is significantly associated with a positive response to immunotherapy (Boll et al., 2025). In order to test if hydrophobicity and positive charge could affect the level of presentation of the peptides, we predicted strong MHC I binders in the protein extensions using MHCflurry (version 2.0.6) (O’Donnell et al. 2018). Overall, we identified 4,369 possible strong binders (size 9 amino acids). We then divided the extensions between those containing at least one predicted strong binder (pMHC) and those not containing it (No pMHC). Out of the 2,808 unique extensions, 1,689 contained at least one predicted strong binder (pMHC).

The chances of containing a predicted MHC I binder will increase with the length of the protein extensions. Consistently, we found that ‘pMHC’ extensions tended to be significantly longer than ‘No pMHC’ ones (Wilcoxon Mann Whitney test p-value = 2.23e-194)(**Figure 2D**). Interestingly, we also found that ‘pMHC’ extensions were clearly more hydrophobic, and had a higher IP, than ‘No pMHC’ ones (Wilcoxon Mann Whitney test p-value = 3.52e-33 and p-value = 2.76e-15, respectively)(**Figures 2E and 2F**). This shows that the same compositional biases that characterize the extensions (high hydrophobicity and positive charge) also favor the presentation of peptides by MHC I molecules. This reinforces the idea that stop-loss mutations can increase the immunogenicity of the tumor by generating “foreign” peptides that are presented by MHC I molecules.

### Enrichment of stop-loss mutations in cancer-associated genes

If stop-loss mutations are associated with tumor-promoting effects, they should be enriched in cancer-associated genes. Examination of a set of 3,347 cancer-associated genes from the Network of Cancer Genes (NCGs)(Dressler et al. 2022) identified 610 stop-loss mutated genes (18.22%). This was a significantly higher fraction than in the case of non-NCGs (2,639 out of 19,884, or 13.27%)(Fisher test p-value 1.381e-13). The relative increase of stop-loss mutations in NCGs *versus* non-NCGs was 37%. These results were also supported by a KEGG cellular pathway enrichment analysis, which revealed that genes in cancer pathways are significantly overrepresented among those with stop-loss mutations (**Figure S2**)(**Table S5**).

In order to examine if there were differences between tumor suppressor genes (TSGs) and oncogenes we analyzed separated gene lists from the COSMIC database (Tate et al. 2019). We found that both types of genes had a very comparable frequency of stop-loss mutations: 71 stop-loss mutated genes in 320 TSGs (22.19%) and 67 stop-loss mutated genes in 319 oncogenes (21%)(Fisher exact test, p-value = 0.7732). Therefore, although stop-loss mutations have been associated with the formation of degrons in tumor suppressor genes (Ghosh et al. 2024), the data suggests that they could also affect the functions of oncogenes. Both classes of cancer genes had similar hydrophobicity and IP distributions (**Figure S3**)(**Table S6**).

### Genes with recurrent stop-loss mutations

Mutation recurrence is a hallmark of cancer-driver mutations (Bailey et al. 2018). We identified 412 genes that had stop-loss in more than one patient (10.97 % of the genes with stop loss mutations), including 61 recurrent mutations found in three or more patients (**Table S6**). The two genes with the highest number of independently observed stop-loss mutations were *PTMA* (14 patients) and *PCDH9* (8 patients)(**Table 1**)(**Table S7**), both with well-studied roles in cancer. Another remarkable case was *SOX9*, recurrently mutated in six different patients.

**Table 1.**
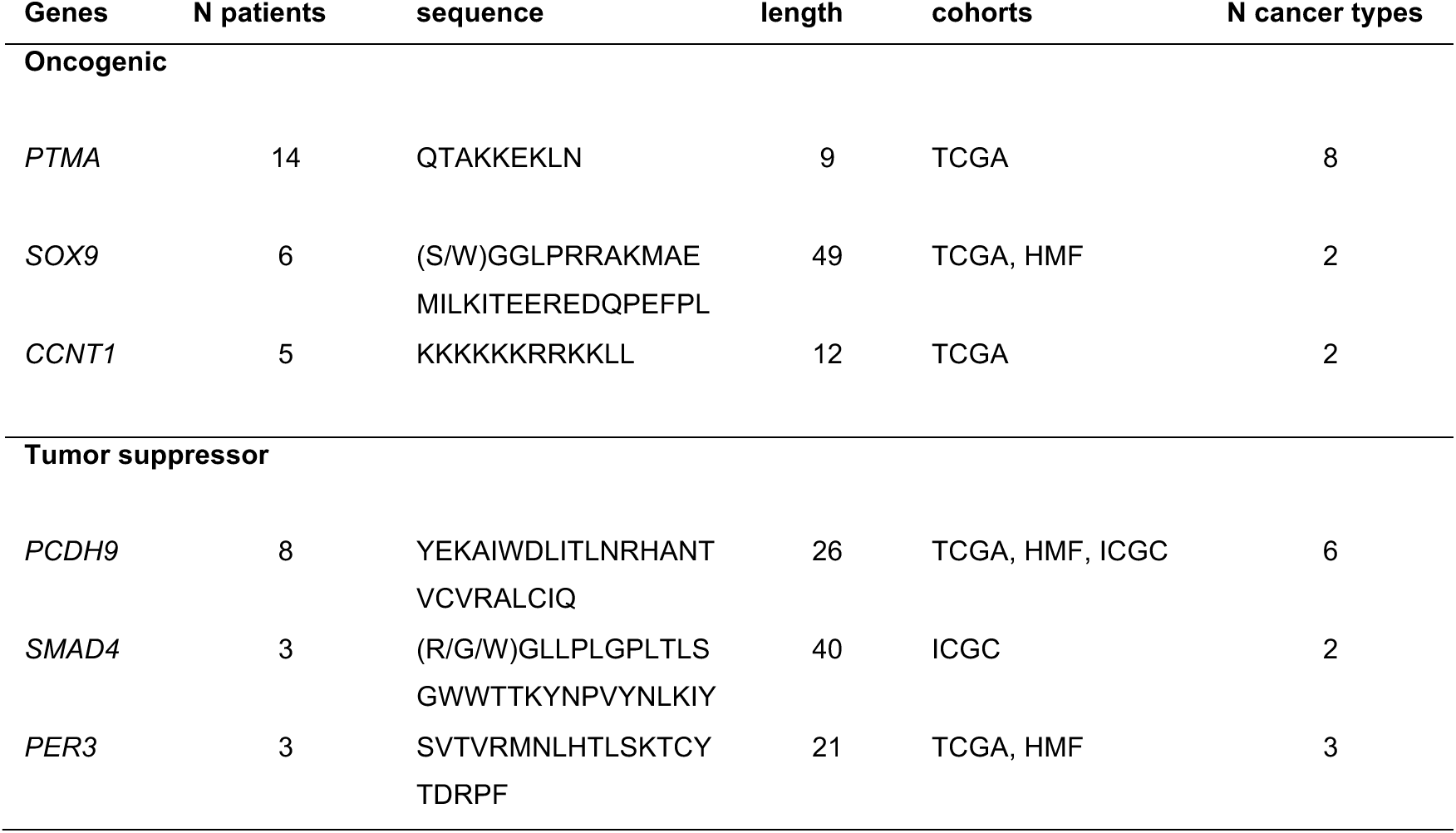
Stop-loss mutated genes in cancer genes. Examples of recurrently mutated genes. Genes are divided into those that have potential tumorigenic effects (Oncogenic) and those that have been associated with suppression of tumor growth (Tumor suppressor). Information on the protein extension sequence, its length, the number of patients with the mutation, and the cohorts/cancer types in which it has been identified, is provided.

*PTMA* encodes the oncoprotein prothymosin alpha (proTα), which is expressed at high levels in different types of cancer (Lin et al. 2015; Kumar et al. 2022). In all fourteen cases, we observed the same T to C mutation resulting in a glutamine. The mutation generated an extension of 9 amino acids with a marked positively charged character (IP 10.51). The second gene with the highest number of stop-loss occurrences (8 cases) was *PCDH9*. This gene has been shown to act as a tumor suppressor gene. For example, in hepatocellular carcinoma, it has been shown to inhibit epithelial–mesenchymal transition and cell migration by activating GSK-3β (Zhu et al. 2014). The mutation was observed in six different types of solid tumors, and the resulting protein extension was 26 amino acids long. *SOX9*, a developmental gene, was also highly recurrently mutated (6 cases). The gene has been shown to promote stemness and inhibit senescence during cancer progression (Matheu et al. 2012; Larsimont et al. 2015). Remarkably, five of the six identified mutations occurred in breast cancer. In this case the protein extension was remarkably long, 49 amino acids, and it contained five putative strong MHC I binders for the common HLA-A02:01 allele (Table S1).

Another example of a recurrently mutated cancer-related gene was *CCNT1.* This gene encodes a cyclin that, together with CDK9, forms the transcription elongation complex P-TEFb. This complex plays an active role in the maintenance of the malignant cell phenotype (Shen et al. 2022). The stop-loss generated an extension of 12 amino acids with a strong positive charge. We also detected recurrent stop-loss mutations in the tumor suppressor gene *SMAD4*, in line with previous studies (Dhamija et al. 2020; Bauer et al. 2023). The gene *PER3*, which controls the circadian rhythm and has been shown to have tumor suppressor activity in multiple cancer types (Tang et al. 2018; Li et al. 2025), displayed stop-loss mutations in 3 different patient samples.

### High expression of *PTMA* in cancer samples

To gain further insights into the effects of stop-loss mutations in proteins with an oncogenic function, we decided to focus on the most recurrently mutated protein, prothymosin alpha (proTα), encoded by *PTMA*. There were 7 cases of testicular germ cell cancer together with individual cases in colorectal cancer, cervical squamous cell carcinoma, esophageal carcinoma, kidney renal clear cell carcinoma, liver hepatocellular carcinoma, sarcoma and uterine corpus endometrial carcinoma. Moreover, the same mutation was found in two cancer patients in the nonstop database (Dhamija et al. 2020), in testicular germ cell cancer and ganglioblastoma (https://nonstopdb.dkfz.de/, accessed 31^st^ of Oct 2024).

The proTα protein is composed of 111 amino acids and a large proportion of the residues have a negative charge (39 glutamic (E) and 18 aspartic acids (D))(**Figure 3A)**. A nuclear transport motif, TKKQKT, has been identified at the protein C-terminus (Manrow et al. 1991). The protein can have an oncogenic effect and modulate the immune system (Samara et al. 2016). Intracellularly, it affects proliferation and cell survival (Enkemann et al. 2000) and prevents apoptosis by inhibiting apoptosome formation (Jiang et al. 2003; Malicet et al. 2006). In the nucleus, the protein promotes chromatin decondensation by histone binding of the central domain (Segade and Gómez-Márquez 1999). PTMA-induced chromatin decondensation around damaged DNA sites, facilitates the local accessibility to the DNA repair machinery (McKnight et al. 2025). PTMA decreases the efficacy of chemotherapeutic agents such as cisplatin (Lin et al. 2016).

**Figure 3:**
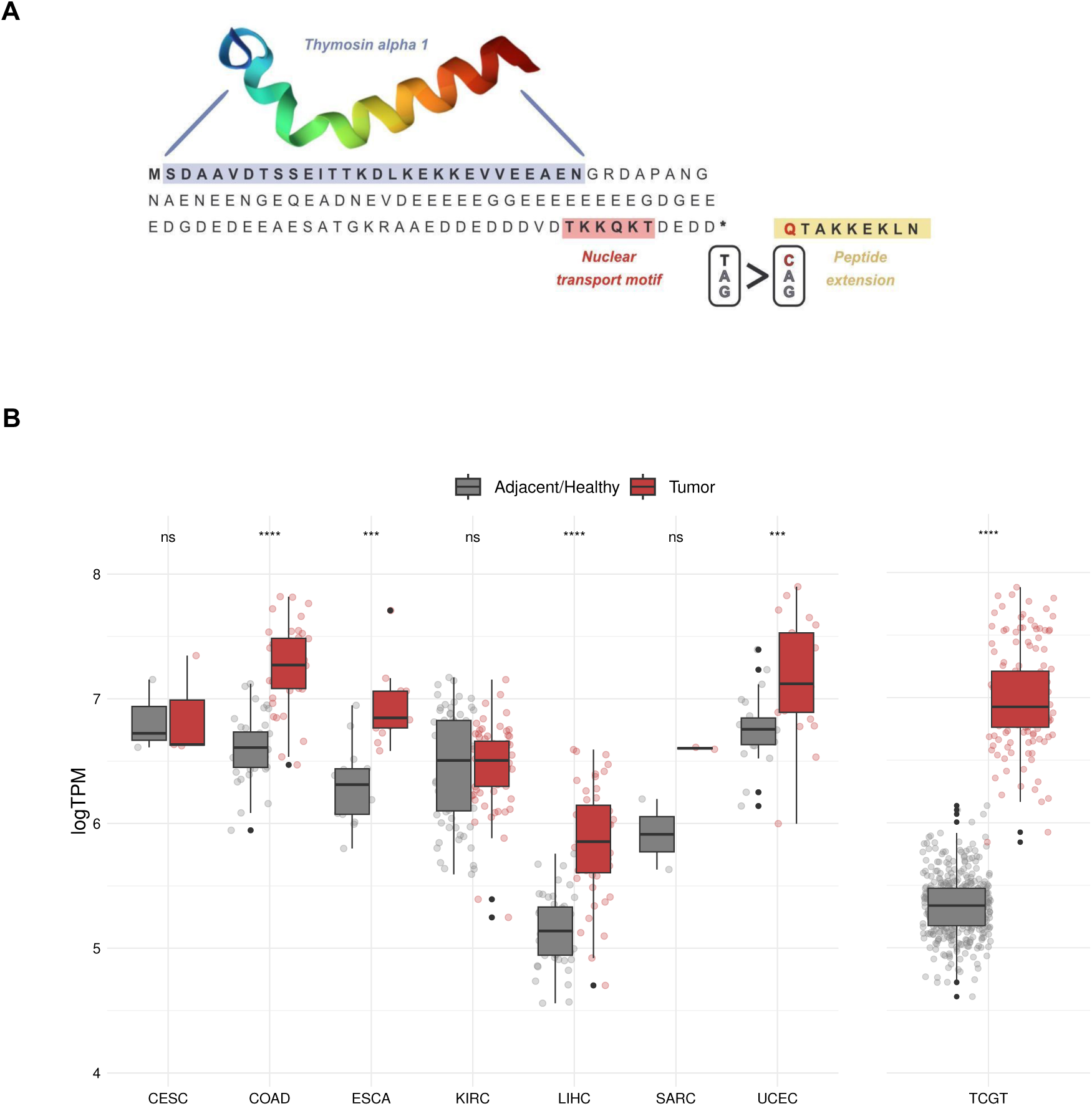
Stop-loss mutations in prothymosin alpha (PTMA) and gene expression in cancer. **A.** Sequence of the PTMA protein with the C-terminal extension generated by the T->C mutation in the stop. Proteolytic cleavage of PTMA generates thymosin alpha 1 (position 2-29). The 3D structure of the thymosin alpha is taken from UniProt P06454. **B.** The expression levels of PTMA tend to be higher in cancer tissues versus controls. Transcript abundance is expressed as logTPM (Transcripts per Million in logarithmic scale). Comparison of the PTMA gene expression values for matched cancer/control data, for different cancer types. CESC: cervical squamous cell carcinoma and endocervical adenocarcinoma; COAD: colon adenocarcinoma; ESCA: esophageal carcinoma; KIRC: kidney renal clear cell carcinoma; LIHC: liver hepatocellular carcinoma; SARC: sarcoma; UCEC: uterine corpus endometrial; TGCT: testicular germ cell tumor. For TGCT, healthy testis expression from the GTEx project is shown compared to the expression in tumor tissue, as no paired data was available. Each dot represents a patient. The line in the box plot indicates the median value. The values were compared using a Wilcoxon test. ns: no significant; ***: p-value < 10e3; ****: p-value <10e4.

Proteolytic cleavage of the N-terminal part of the proTα protein generates the 28 amino acid peptide thymosin alpha 1, which has been shown to enhance the activity of T cells and natural killer (NK) cells (Garaci et al. 2012; Wang et al. 2023). Because of its immunostimulatory effects, this peptide has been used therapeutically to treat cancer (Moody 2007; Ioannou et al. 2012). Therefore, in contrast to the proliferative effects of proTα protein, the activity of the thymosin alpha 1 can protect against cancer by activating the immune system.

Analysis of *PTMA* gene expression in different tissues from GTEx (The GTEx Consortium 2013) supported ubiquitous expression of the gene. High expression levels were also observed in EBV-transformed lymphocytes displaying very high *PTMA* mRNA abundance (**Figure S4**)(**Table S9**). In accordance with early findings of an increased expression level for *PTMA* in malignant tissue compared to healthy tissue (Kobayashi et al. 2006; Tsai et al. 2009), we observed that *PTMA* tended to be overexpressed in cancer tissues when compared to matched controls (**Figure 3B**). The trend was particularly strong in tumor tissues in which *PTMA* stop-loss mutations had been observed, such as testicular germ cell tumor (TGCT), liver hepatocellular carcinoma (LIHC), and colon adenocarcinoma (COAD). In TGTC samples with mutation and expression data available, the *PTMA* stop-loss mutation did not show any clear association with changes in transcript levels (**Figure S5).**

### The stop-loss mutation in *PTMA* alters the cleavage of the N-terminal part of the protein

To determine whether the stop-loss mutation in *PTMA* leads to an altered protein function, we cloned the open reading frames (ORF) of wild-type (WT) PTMA and stop-loss extended forms in an expression vector. JHH6, HuH7 and HeLa cells were selected as representatives for liver and cervical cancers, where stop-loss mutants of PTMA have been described. We determined that cell proliferation is similar in JHH6 transiently (**Figure 4A, Table S11**) or stably (**Figure 4B, Table S12**) expressing WT or extended PTMA and is higher compared to control cells, as expected. Similarly, control cells were more sensitive to cisplatin than JHH6 cells stably expressing PTMA (**Figure 4C, Table S13**). However, there are no significant differences in the resistance to this therapy between cells expressing WT or extended PTMA.

**Figure 4:**
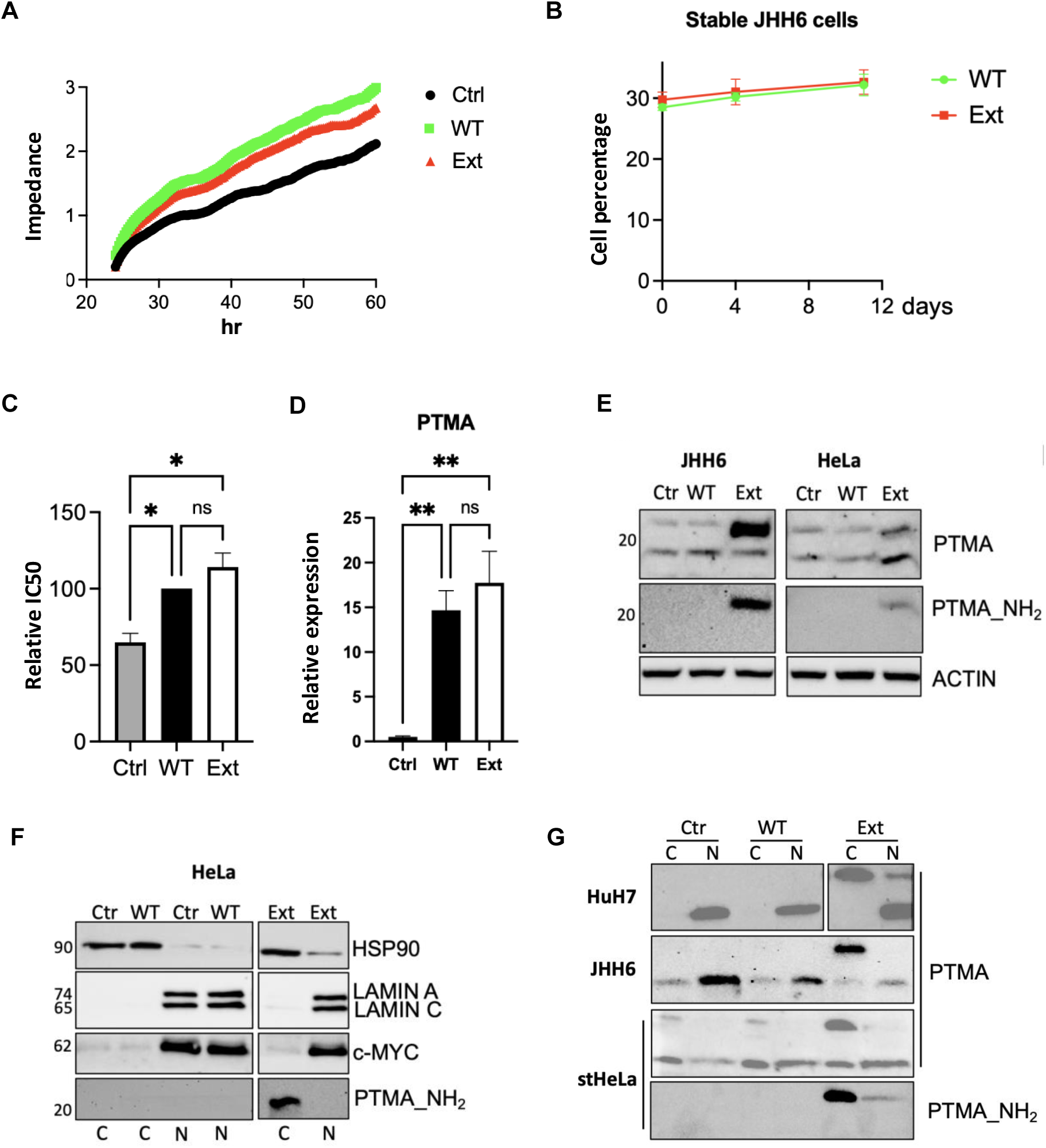
Functional analysis of stop-loss mutation in PTMA. **A-B.** Cell growth was evaluated in JHH6 cells transfected with control plasmids (Ctrl) or expressing wild-type (WT) or extended (Ext) PTMA or in JHH6 cells stably expressing these plasmids. **C.** JHH6 stably expressing the plasmids were incubated with cisplatin and the IC50 was calculated and plotted as a percentage. **D.** JHH6 liver cancer cells were transfected as in A and PTMA levels were evaluated by quantitative RT-PCR. RPLP0 was used as a normalizer. **E-G.** The indicated proteins were visualized by Western blot in extracts from cells (JHH6, HeLa or HuH7) transfected as indicated in A. Extracts were total (E) or cytoplasmic or nuclear fractions (F and G). PTMA_NH2 corresponds to an antibody against the N-terminal part of the protein. HeLa cells stably expressing the plasmids (stHeLa) were also evaluated. All experiments were performed at least twice, and the average result is shown with standard deviation (B, C, D) or a representative result is shown (E to G). ANOVA results are shown with Šídák’s multiple comparison test. ns: non-significant. *p-value < 0.05. ** p-value < 0.01.

Transfection of these cells with plasmids expressing the WT or extended forms of PTMA shows similar levels of *PTMA* mRNA (**Figure 4D, Table S14**) and PTMA protein of the expected size (around 18 kDa)(**Fig. 4E**). However, the cells transfected with the extended version of PTMA show an additional protein of around 21 kDa. Given that the size of the thymosin alpha 1 peptide that is cleaved from the NH2 part of PTMA is 3 kDa, we considered that the large PTMA could correspond to the uncleaved protein (18 kDa + 3 kDa). This was verified with a PTMA antibody that only recognizes the NH2-termini of PTMA (**Figure 4E**). Since the extension of the stop-loss mutant is close to the nuclear localization signal and could affect its nuclear targeting, we fractionated transfected cells and determined that the large PTMA protein accumulates in the cytoplasmic fraction (**Figure 4F**). Instead, the WT protein is found in the nuclear fraction in all evaluated cells (**Figure 4G**). This was also the case in cells that stably overexpressed the WT and extended versions of PTMA (stHela). While altered cleavage or localization of the extended PTMA did not correlate with differential cell growth or resistance to therapy, decreased levels of thymosin alpha 1 peptide in the stop-loss PTMA mutant could potentially cause lower immunogenicity and higher tumor progression.

## DISCUSSION

Studies on mutated genes in cancer have mostly focused on missense mutations; in contrast, stop-loss mutations have been barely investigated. Here we combined data from 20,801 cancer patients to undertake a genome-wide survey of stop-loss mutations. We found that cancer-related genes were significantly enriched in stop-loss mutations, and that the three genes with the highest number of stop-loss mutation recurrences had oncogenic or tumor suppressor functions. These findings provide strong evidence that stop-loss mutations can play roles in the development of cancer.

Mutation recurrence is well documented in cancer-driver genes (Tamborero et al. 2013; Bailey et al. 2018). In the case of oncogenes, the gene can increase its activity by increasing gene expression levels or by specific mutations that make them hyperactive. For example, 90% of the B-raf proto-oncogene (*BRAF*) mutations are a substitution of a V by an E in position 600, which causes continuous cell proliferation (Davies et al. 2002). In contrast, tumor suppressor genes, such as P53, tend to accumulate protein-inactivating mutations. The effects of stop-loss mutations on protein functions are only starting to be investigated. It has been shown that stop-loss mutations can generate hydrophobic degron motifs that promote the degradation of the protein (Ghosh et al. 2024); these effects would have been reported in *SMAD4* and *VHL* (Dhamija et al. 2020; Pal et al. 2025). A recent massively parallel analysis has shown that the polypeptides translated from non-coding sequences often have hydrophobic C-terminal tails, leading to degradation or membrane targeting of the proteins (Kesner et al. 2023).

Here we quantified the differences in hydrophobicity and IP in the protein extensions generated by stop-loss mutations when compared to standard proteins. We found that, in general, protein extensions were more hydrophobic and positively charged than canonical proteins. Interestingly, it has been reported that hydrophobic sequences are more likely to be loaded onto the MHC-I complex by proteasomal degradation than other types of sequences (Seong and Matzinger 2004). In our study, MHC I peptide binders predictions also supported that high hydrophobicity increases the likelihood of MHC I binding. Additionally, a high isoelectric point had the same effect, further supporting the idea that stop-loss mutations could make the cells more immunogenic. These biases could also result in the generation of novel protein functional motifs. For example, clusters of lysine and arginine could generate novel nuclear localization signals, influencing the subcellular localization of the protein. Positively charged sequences might also favor the interactions with nucleic acid sequences, which have a negative charge, or with protein regions which are rich in acidic amino acids.

We discovered several cancer related genes with recurrent stop-loss mutations, pointing to tumorigenic effects linked to these mutations. The one with the highest number of occurrences was *PTMA,* which encodes the protein prothymosin alpha (proTα), known to promote cell proliferation. The protein C-terminal extension, containing 9 amino acids, was positively charged. We found that, in accordance with the oncogenic nature of this gene, *PTMA* overexpression led to increased cell proliferation and resistance to therapy. However, these effects were similar in cells expressing WT PTMA or the stop-loss extended mutant. However, the mutant showed a higher accumulation in cytoplasmic fractions. Given that PTMA nuclear localization signal is close to the carboxi-termini, where the extension is located, we speculate that an interference between these sequences may affect nuclear translocation. The extension could also potentially affect post-translational modifications in the carboxi-part, such as phosphorylations, sumoylations and ubiquitinations.

Although the oncogenic role of *PTMA* is widely accepted, *PTMA* can also play pro-inflammatory roles. Extracellularly it enhances the immune system by promoting dendritic cell maturation, binding to Toll-like receptor 4 on macrophages and neutrophils (Romani et al. 2004), and increasing T lymphocyte activity in combination with interleukin-2 (Voutsas et al. 2000). ProTα is also the precursor of Thymosin alpha 1 by cleavage at the N-terminal by asparaginyl endopeptidase (Chen et al. 2006). This peptide has been associated with immune regulation by enhancing T cell differentiation, antibody responses and chemokine production, and it has shown to cause a decrease of tumor growth in different animal models (Beuth et al. 2000; Chen et al. 2006; Moody 2007; Sungarian et al. 2009). Our results indicate that cells expressing the stop-loss extended *PTMA* showed high levels of a large protein recognized by thymosin alpha 1 antibodies. This suggests that the carboxy-terminal extension may affect the amino-terminal cleavage in a direct (by affecting protein structure and peptidase cleavage) or indirect manner (by altering localization). While the mechanism for this is unclear, a decreased cleavage and less accumulation of thymosin alpha 1 in stop-loss mutant tumor cells should lead to lower immunogenicity, favoring tumor growth.

Our results provide evidence that stop-loss mutations accumulate in cancer cells, and that they do so more often in genes that are known to be related to cancer than in other types of genes. By modulating the stability and activity of the proteins, and creating novel immunogenic peptides, these mutations can shape tumor behavior and influence therapeutic responses. Future studies in which mutation data can be studied in conjunction with immunopeptidomics data for a large number of tumors will shed new light into how stop-loss mutations can increase the recognition of the tumor by the immune system.

## METHODS

### Somatic mutation information

We downloaded mutation tables from the NCI Cancer Genome Characterization initiative (CGCI, phs000235), available through dbGaP (CGCI-BLGSP (phs000527), CGCI-HTMCP-CC (phs000528), CGCI-HTMCP-DLBCL (phs000529), CGCI-HTMCP-LC (phs000235)). We also downloaded the publicly available somatic mutation data of the 33 TCGA cohorts from the MC3 pipeline (https://gdc.cancer.gov/about-data/publications/mc3-2017, accessed: 8th of October 2024), as well as the final_consensus_passonly.snv_mnv_indel maf file of the ICGC/PCAWG cohort following the instructions on their website (https://docs.icgc-argo.org/docs/data-access/icgc-25k-data#open-release-data---object-bucket-details, accessed: 22nd of April 2025). We also gained access to somatic variant data from tumors from the Stitching Hartwig Medical Foundation via a License Agreement. Annotated mutation tables of seven additional datasets were downloaded from the corresponding publications: CPTAC-3 (Li et al. 2023), HCMI- CMDC (phs001486), MMRF- COMMPASS (Keats et al. 2013), WCDT- MCRPC (Zhao et al. 2020) and other studies (Bassani-Sternberg et al. 2016; Löffler et al. 2019; Roper et al. 2020). Whenever necessary, we lifted the data from the GRCh37 to the GRCh38 version of the human genome. When available, all stop-loss mutations were filtered for a population-wide allele frequency of < 5% (gnomAD), sample depth ≥ 30X, and alternative allele depth ≥ 3X. In cases in which there were multiple tumor samples per patient we only kept the set of non-redundant mutations.

We also processed raw whole exome sequencing data from paired germline and tumor samples of six cohorts (Mariathasan et al. 2018; Boll et al. 2023; Miao et al. 2018; Snyder et al. 2017; Chong et al. 2020; Kraemer et al. 2023). Sequencing reads were trimmed with cutadapt (v4.1) and quality-checked using FastQC (version 0.11.7). Reads were aligned to GRCh38 using BWA (version 0.7.17) and, in cases of several fastq files per patient, merged with GATK MergeSamFiles. Base recalibration and duplicate marking were done with GATK BaseRecalibrator. Contamination was estimated with GATK CalculateContamination, and coverage was assessed using Qualimap (version 2.2.1). Alignment metrics were collected with GATK CollectAlignmentSummaryMetrics. Mutations were called using GATK Mutect2, Strelka2 (v2.9.10), and VarScan2 (version 2.4.4), with SAMtools (version 1.12) mpileup providing input for VarScan2. Mutect2 used the germline-resource file somatic-hg38_af-only492 gnomad.hg38.vcf.gz, and filtering was done with GATK FilterMutectCalls. Only PASS mutations were retained. An ensemble mutation file, requiring detection by at least two callers, was generated using bcbio-variation-ensemble (version 0.2.2.6) and annotated with VEP (version 104). We generated maf files using vcf2maf (version 0.1.16). Finally, mutations were filtered for a population-wide allele frequency of < 5% (gnomAD), sample depth ≥ 30X, and alternative allele depth ≥ 3X. Detailed information from stop-loss mutations is available from Table S1.

### Lists of cancer genes

The cancer-related genes were taken from the Network of Cancer Genes including genes found in at least two of the analysed screens (http://network-cancer-genes.org/, last accessed February 27^th^ 2026)(Dressler et al. 2022).

The gene sets of tested tumor suppressors and oncogenes were obtained from COSMIC (https://cancer.sanger.ac.uk/cosmic/census?tier=2, last accessed September 11th, 2023)(Tate et al. 2019).

### Identification of stop-loss mutations and protein extensions

We extracted all the mutations affecting stop codons from the somatic variant tables to build a catalogue of nucleotide mutation frequencies at this position. The same was applied to the missense mutations. We computed the amino acid that would result from the mutation using the standard human genetic code and information from the 3’UTR sequences of the gene annotations from GRCh38, Ensembl release 112. We retrieved a total of 3,148 extensions, of which 2,808 were unique. The code for the extension prediction can be accessed at https://github.com/justalilibit/stop-loss-muts_in_canonical. Detailed information of the protein C-terminal extensions is available from Table S1.

### Identification of MHC I peptide binders derived from stop-loss mutations

The binding affinity of all possible peptides of size 9 amino acids in the protein extensions to the most common MHC I molecules was predicted using MHCflurry (version 2.0.6)(O’Donnell et al. 2018), applying a stringent IC50 threshold of 50 nM. A list of all predicted strong binders can be found in Table S1.

### Hydrophobicity and isoelectric point computations

We downloaded the database of canonical proteins from UniProt and SwissProt to compare the properties of peptide extensions from stop-loss mutations with those of canonical sequences. We predicted intronic open reading frames (iORFs) in human introns (Genome assembly hg38) using orfipy (version 0.0.4), with a minimum peptide length of 9 amino acids.

We used the R package ‘Peptides’ (version 2.4.6) to calculate the GRAVY hydrophobicity index with the “KyteDoolittle” scale, and the isoelectric point with the EMBOSS pK-scale.

### Gene expression data

We downloaded gene expression values directly from the TCGA (Weinstein et al. 2013) and GTEx projects (The GTEx Consortium 2013). Expression values were measured as transcripts per Million or TPM. Whenever possible, we used matched cancer/control gene expression data from TCGA. For testicular cancer (TCGT), TCGA does not contain adjacent data. Instead, we used healthy testis expression data from the GTEx project version 10.

### Cell lines

Human HeLa (cervical carcinoma) and HuH7 (hepatocellular carcinoma, HCC) cell lines were cultured in Advanced DMEM (Gibco, Life Technologies) supplemented with 10% fetal bovine serum (FBS), glutamine and antibiotics. JHH6 (HCC) cells were maintained in William’s E medium supplemented with 10% FBS, glutamine and antibiotics. All cell lines were routinely tested for mycoplasma contamination using the MycoAlert Mycoplasma Detection Kit (Lonza, #LT07–318). Cells were maintained at 37 °C in a humidified atmosphere containing 5% CO₂.

### DNA cloning

PTMA WT and extended ORF sequences with a Kozak in the initial ATG were synthesized by GeneScript with SfiI and Xba I neighboring restriction sites and were cloned into the same sites of the sleeping beauty backbone plasmid pSB-GFP-PurCherry (Rovira et al. 2023). Positive clones were verified by Sanger sequencing and named pSB-PTMA-WT or pSB-PTMA-Ext.

### Transient transfection and generation of stable cell lines

HeLa, Huh7, and JHH6 cells were seeded at a density of 200,000 cells per well in 6-well plates and transfected the following day using Lipofectamine 2000 (Thermo Fisher Scientific), according to the manufacturer’s instructions. A total of 1 µg of pSB-PTMA-WT or pSB-PTMA-Ext and 2 µL of Lipofectamine 2000 were used per well.

For the generation of stable cell lines, cells were co-transfected with pSB-PTMA-WT or pSB-PTMA-Ext containing an mCherry reporter, together with a plasmid expressing the Sleeping Beauty transposase. At 72h post-transfection, cells were selected with puromycin (1.75 and 1 µg/mL for HeLa and JHH6, respectively) until non-transfected control cells were eliminated. Puromycin-resistant cells were expanded, and the proportion of transgene-positive cells was determined by flow cytometry based on mCherry fluorescence.

### Evaluation of RNA expression by RT-qPCR

Total RNA was extracted with a Maxwell® RSC simpleRNA Tissue Kit (Promega) following the manufacturer’s protocol. RNAs were quantified by measuring absorbance at 260 nm using a NanoDrop™ 1000 (Thermo Fisher Scientific). Complementary DNA (cDNA) was obtained by M-MLV-based (Invitrogen) reverse transcription of one µg total RNA and was evaluated by RT-qPCR in a CFX96 Real-Time system (Bio-Rad). RT-qPCRs were performed using IQ SYBR Green Supermix (Bio-Rad), 0.4 µM primers (forward and reverse), and two µL cDNA in a total reaction volume of ten µL. Primers were designed using the Primer3 platform and synthesized by Thermo Scientific. Primer sequences were PTMAF: GGCTGACAATGAGGTAGACGAAG, PTMAR: GTAGCTGACTCAGCTTCCTCATC; RPLP0F: AGCCCAGAACACTGGTCTC and RPLP0R: ACTCAGGATTTCAATGGTGCC.

### Protein extraction, cell fractionation and Western blot analysis

For total protein extraction, cells were lysed in RIPA buffer containing NaCl, Tris-HCl, deoxycholate, SDS, Triton X-100, and a cocktail of phosphatase (1 mM sodium orthovanadate, 10 mM sodium fluoride, 100 mM β-glycerophosphate) and protease inhibitors (Roche). Lysates were sonicated, clarified by centrifugation (16,000 × g, 30 min, 4 °C), and protein extracts were analyzed by Western blotting. PTMA was detected using anti-PTMA antibodies Thermo Fisher Scientific #PA5-71580 (recognizing the region 1-33 amino acids) and Abcam #ab247074, both 1:1000, with B-actin (Sigma-Aldrich, #A2066, 1:1000) as a loading control.

For subcellular localization analyses, parental, transiently transfected, and stable HeLa cells expressing wild-type or extended PTMA were subjected to cellular fractionation followed by immunoblotting. Cells were harvested, washed with ice-cold PBS, and resuspended in hypotonic buffer (10 mM HEPES pH 7.9, 1.5 mM MgCl₂, 10 mM KCl, 0.5 mM DTT, 0.5 mM PMSF, protease inhibitors), incubated on ice for 10 min, supplemented with 0.5% NP-40, and lysed by gentle pipetting. After centrifugation (800 × g, 10 min, 4 °C), the supernatant was collected as the cytoplasmic fraction. Nuclear pellets were extracted in hypertonic buffer (20 mM HEPES pH 7.9, 420 mM NaCl, 0.2 mM EDTA, 20% glycerol, 0.5 mM DTT, 0.5 mM PMSF) for 30 min at 4 °C and clarified by centrifugation (16,000 × g, 15 min, 4 °C). Fraction purity was validated by immunoblotting using Hsp90 (Cell Signaling Technology, #4874, 1:1000) as cytoplasmic markers, and lamin A/C (Cell Signaling Technology, #2032, 1:1000) and c-myc (Cell Signaling Technology, #5605, 1:1000) as nuclear markers.

### Competitive co-culture assay monitored by flow cytometry

Competitive co-culture assays were performed using human JHH6 parental cells (mCherry-negative) mixed with stable cell lines expressing *wild-type* (WT) PTMA or the extended PTMA variant, all co-expressing mCherry. Mixed cell populations were seeded and analyzed at day 0 and after 4, 7, and 11 days post-seeding. Cells were harvested and the proportion of mCherry-positive cells was quantified by flow cytometry using a CytoFLEX 4LS (Beckman Coulter) as a readout of proliferative advantage.

### Real-time cell proliferation assay (xCELLigence RTCA)

Cell proliferation and growth kinetics were monitored in real time using the Agilent xCELLigence Real-Time Cell Analysis (RTCA) system, which measures electrical impedance as an indicator of cell number. Background impedance was first measured by adding 100 µL of complete culture medium to each well of an E-Plate 16. Cells were harvested and counted using an automated cell counter, and 6,000 were seeded in a final volume of 200 µL per well. Plates were inserted into the RTCA station inside a humidified incubator (37 °C, 5% CO₂) and impedance measurements were recorded automatically every 15 min for 96 h. Data were analyzed using RTCA software and exported for downstream statistical analysis and visualization. Experiments were performed with four technical replicates per condition and repeated independently three times.

### Cell viability and IC₅₀ determination by MTS assay

Cell viability and cisplatin IC₅₀ values were determined using a colorimetric MTS assay (Promega CellTiter 96®). HeLa and JHH6 cells were seeded in 96-well plates at a density empirically determined to avoid reaching confluence at the end of the 72 h experiment. Cells were allowed to attach overnight before treatment. Cisplatin was added in a serial dilution covering a concentration range from 80 µM to 0.3 µM in complete culture medium. Control wells received vehicle only. Each condition was plated in three technical replicates. Cells were incubated with the drug for 72 h under standard culture conditions (37 °C, 5% CO₂). At the end of the treatment period, MTS reagent was added to each well according to the manufacturer’s instructions and plates were incubated at 37 °C until sufficient color development was achieved (typically 1 h). Absorbance was measured at 490 nm using a microplate reader. Background signal from medium-only wells was subtracted, and cell viability was calculated relative to untreated controls. Dose–response curves and absolute IC₅₀ values were determined by nonlinear regression analysis using GraphPad Prism 8. Experiments were performed in at least three independent biological replicates. Due to experimental variability observed in IC₅₀ values across independent experiments, IC₅₀ values from each experiment were normalized to the IC₅₀ of the wild-type (WT) control from the same experiment.

### Statistical analysis

Plots were generated using Python (version 3.8.6), R (version 4.1.2), and Rstudio (version 1.4). We used the Fisher exact test to compare proportions between groups, and the Mann Whitney test to compare distributions of continuous variables.

## AUTHOR CONTRIBUTIONS

L.M.B. performed most of the bioinformatics analyses, including variant calling, statistics on stop-loss mutations, identification of protein extensions and prediction of MHC I binders. J.A.M. collected mutation data from the Hartwig database, reanalyzed the data and performed data curation. M.E.C. contributed to the prediction of putative MHC I binders and to the study of *PTMA* gene expression levels. J.P.B contributed to the conceptualization of the study and analysis of the mutation data. L.M.B and M. M.A. conceived the study and designed the main bioinformatics analyses. N.K.K. made constructs for the studies of *PTMA* in cell lines. N.K.K., C.V., S.S.V., E.S and I.A. performed Western blots. J.C.G., N.K.K. and E.S. performed experiments to assess cell growth and resistance to therapy by different methods. P.F. led the experimental part of the study. L.M.B., P.F, M.M.A wrote the manuscript with contributions from all co-authors.

## ACKNOWLEDGEMENTS

This publication and the underlying study have been made possible partly based on data that Hartwig Medical Foundation has made available to the study through the Hartwig Medical Database. We acknowledge funding from Ministerio de Ciencia, Innovación y Universidades grant PID2021-128791OB-I00, PID2024-162742OB-I00, and PID2021-122726NB-I00 funded by MCIN/AEI/10.13039/501100011033 and by “ERDF: A way of making Europe”, by the “European Union”. We also acknowledge funding from Generalitat de Catalunya, grant 2021SGR00042 and the Instituto de Salud Carlos III, which finances Centro de Investigación Biomédica en Red de Enfermedades Hepáticas y Digestivas (CIBEREhd), financed by the EU (NextGenerationEU). Plan de Recuperación Transformación y Resiliencia). L.M.B. received funding from La Caixa INPHINIT doctoral program. N. K. K. was funded by grant number PRE2022-101505 and I. A. from grant number PREP2024-002818 from Ministerio de Ciencia, Innovación y Universidades.

