## Supplementary Figures for "Oncogenes and tumor suppressor genes are enriched in stop-loss mutations generating protein extensions"

| **TAG** | * | **TAA** | * | **TGA** | * |
|---|---|---|---|---|---|
| T**C**G | Ser (S) | T**C**A | Ser (S) | **G**GA | Gly (G) |
| TA**C** | Tyr (T) | TA**C** | Tyr (T) | T**C**A | Ser (S) |
| TA**T** | Tyr (T) | TA**T** | Tyr (T) | TG**C** | Cys (C) |
| T**G**G | Trp (W) | T**T**A | Leu (L) | T**T**A | Leu (L) |
| T**T**G | Leu (L) | **C**AA | Gln (Q) | TG**G** | Trp (W) |
| **C**AG | Gln (Q) | **G**AA | Glu (E) | TG**T** | Trp (W) |
| **G**AG | Glu (E) | **A**AA | Lys (K) | **A**GA | Arg (R) |
| **A**AG | Lys (K) | TA**G** | * | **C**GA | Arg (R) |
| TA**A** | * | T**G**A | * | T**A**A | * |

| hydrophobic | polar | basic | neutral | acidic |
|---|---|---|---|---|

**Figure S1. Possible single nucleotide variants (SNVs) in the stop codons and the resulting amino acid changes.** The majority of stop codon mutations extend the coding sequence. Mutated codons with the encoded amino acid and its chemical properties. Variants that encode another stop codon (*) are marked in grey.

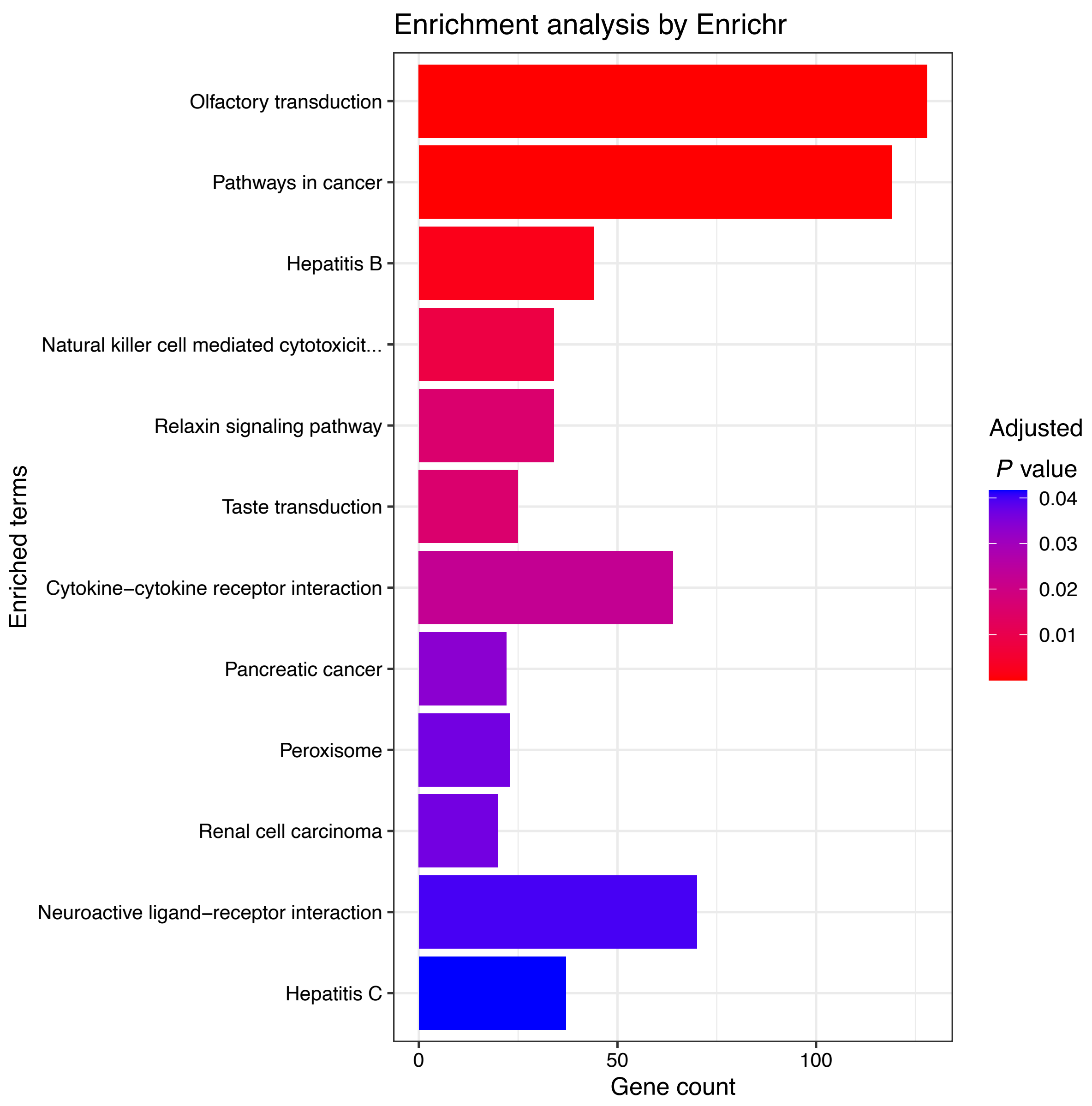


**Figure S2. Gene enrichment analysis of stop-loss mutated genes in KEGG pathways.** Pathway enrichment was performed using Enrichr with the KEGG_2021_Human database. The top 20 enriched pathways are shown, ranked by adjusted p-value. Bar lengths represent the number of stop-loss mutated genes overlapping with each pathway. Missense-mutated genes were used as the background set to control for general mutation bias. Olfactory transduction, pathways in cancer and hepatitis B are the pathways that are significantly overrepresented among stop-loss variants (adj. p-value < 0.05).

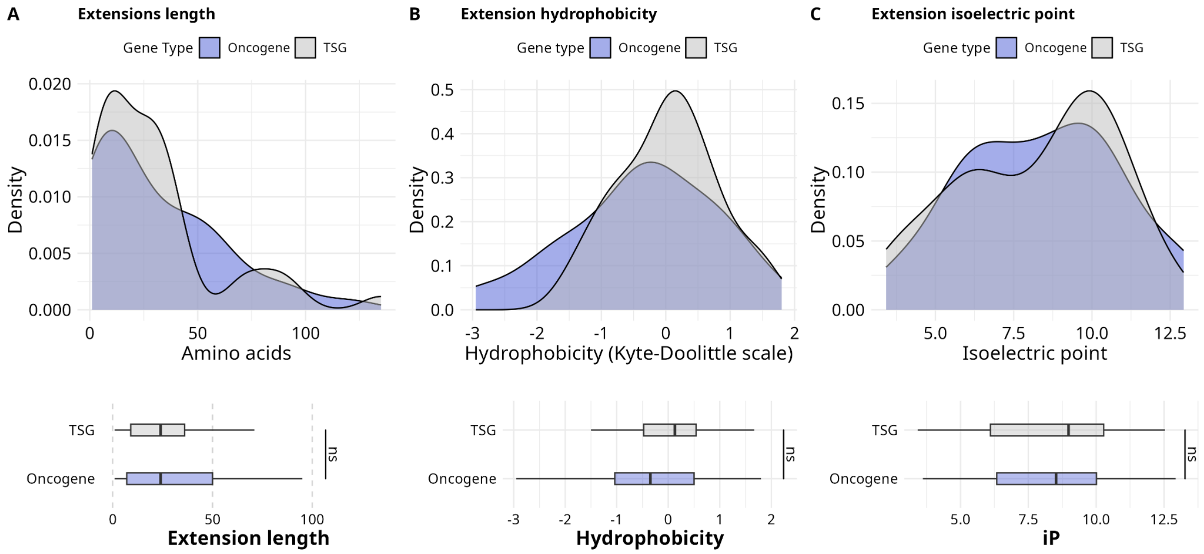


**Figure S3. Properties of protein extensions in tumor supressor genes (N = 42) versus oncogenes (N = 43)**. **A. Protein extension length**. No significant difference in the length of the peptide extension between the two gene sets (p-value = 0.98). **B. Protein extension hydrophobicity**. Stop-loss extensions from tumor suppressor genes show a higher median hydrophobicity but the difference is not significantly different from extensions by oncogene (p-value = 0.15). **C. Protein extension isoelectric point.** The distribution of the isoelectric point of the extensions is not  in the TSG and oncogene group (p-value = 0.94). P-values were obtained with the Wilcoxon-Mann-Withney test.

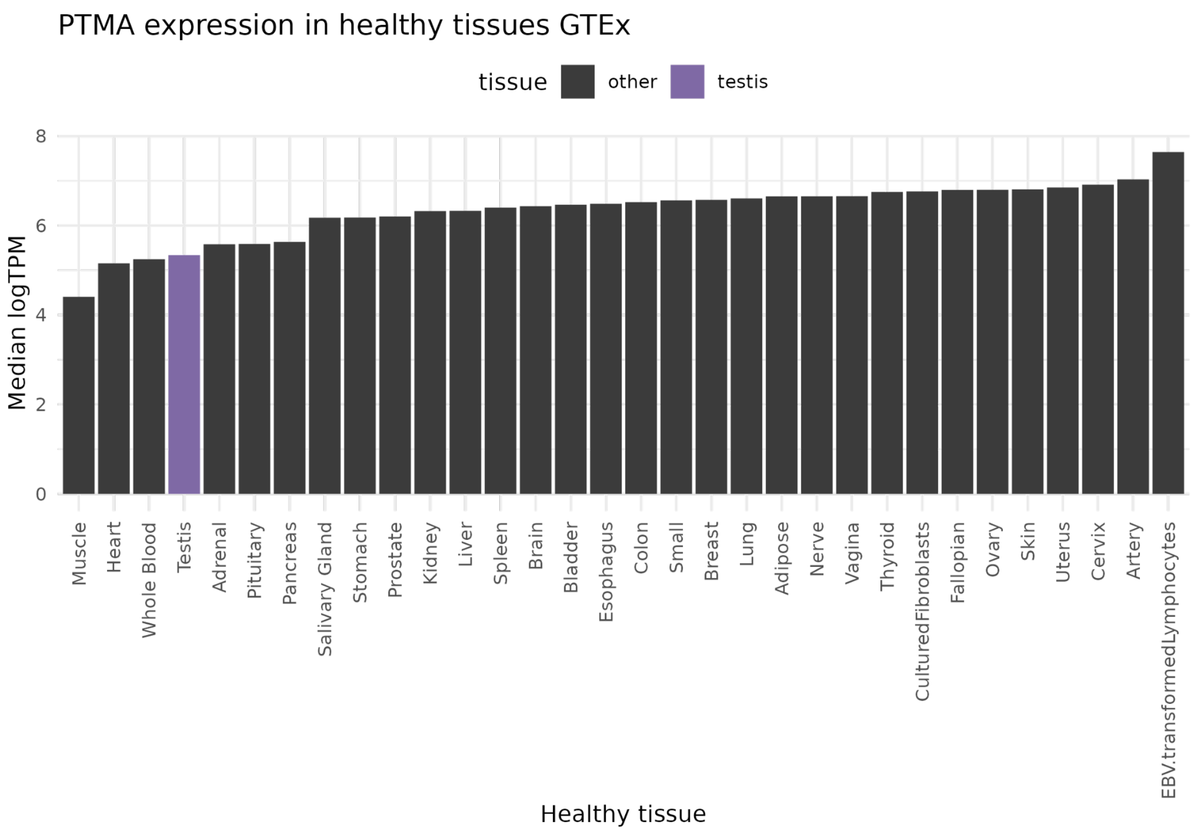


**Figure S4. Expression of the *PTMA* gene in different body tissues**. The median logTPM value in GTEX is shown.

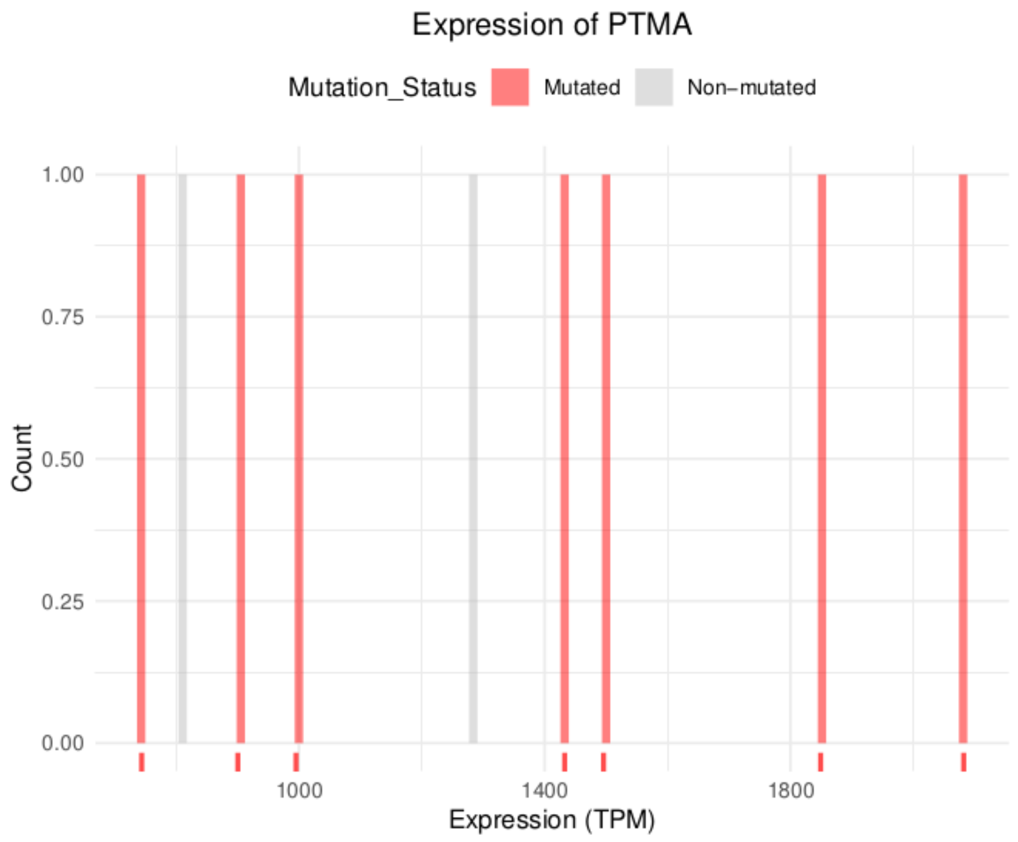


**Figure S5: Expression of *PTMA* in TGCT patients with and without the stop-loss mutation**. Patients with a stop-loss mutation in PTMA are marked red. The expression in mutated samples is not significantly different from the distribution of expression values. Gene expression values are in Transcripts per Million (TPM). Data is from TCGA.
